# pegFinder: A pegRNA designer for CRISPR prime editing

**DOI:** 10.1101/2020.05.06.081612

**Authors:** Ryan D. Chow, Jennifer S. Chen, Johanna Shen, Sidi Chen

**Affiliations:** Department of Genetics, Yale University School of Medicine, New Haven, Connecticut, USA; System Biology Institute, Yale University, West Haven, Connecticut, USA; Center for Cancer Systems Biology, Yale University, West Haven, Connecticut, USA; M.D.-Ph.D. Program, Yale University, West Haven, Connecticut, USA; Department of Laboratory Medicine, Yale University School of Medicine, New Haven, Connecticut, USA; Department of Immunobiology, Yale University School of Medicine, New Haven, Connecticut, USA; Immunobiology Program, Yale University, New Haven, Connecticut, USA; Combined Program in the Biological and Biomedical Sciences, Yale University, New Haven, Connecticut, USA; Yale Comprehensive Cancer Center, Yale University School of Medicine, New Haven, Connecticut, USA; Department of Neurosurgery, Yale University School of Medicine, New Haven, Connecticut, USA; Yale Stem Cell Center, Yale University School of Medicine, New Haven, Connecticut, USA; Yale Liver Center, Yale University School of Medicine, New Haven, Connecticut, USA; Yale Center for Biomedical Data Science, Yale University School of Medicine, New Haven, Connecticut, USA; Center for RNA Science and Medicine, Yale University School of Medicine, New Haven, Connecticut, USA

## Abstract

CRISPR technologies have been widely adopted as powerful tools for targeted genomic manipulation ^1^. Recently, a new CRISPR-based strategy for precision genome editing was developed that enables diverse genomic alterations to be directly written into target sites without requiring double-strand breaks (DSBs) or donor templates ^2^. Termed prime editing, this approach involves two key components: 1) a catalytically impaired Cas9 nickase fused to a reverse transcriptase (PE2), and 2) a multifunctional prime editing guide RNA (pegRNA) that specifies the target site and further acts as a template for reverse transcription (RT). pegRNAs are similar to standard single-guide RNAs (sgRNAs), but additionally have a customizable extension on the 3’ end. The 3’ extension is composed of a RT template that encodes the desired edit and a primer binding site (PBS) that anneals to the target genomic site to prime the RT reaction ^2^. These additional components considerably increase the complexity of pegRNA design compared to standard sgRNAs. While many tools have been developed for identifying candidate sgRNAs in a target DNA sequence ^3–8^, no user-friendly web application currently exists for designing pegRNAs. We therefore developed pegFinder, a streamlined web tool that rapidly designs candidate pegRNAs (**Figure 1**). The pegFinder web portal is freely available at http://pegfinder.sidichenlab.org/ (**Supplementary Figure 1**).

---

In **standalone** mode, pegFinder simply requires two inputs: 1) the wildtype/reference DNA sequence of the target site and its flanking regions, and 2) the edited/desired DNA sequence. For consideration of predicted on-target efficacy scores, the user can optionally include the results from the Broad sgRNA designer (https://portals.broadinstitute.org/gpp/public/analysis-tools/sgrna-design) or CRISPRscan (https://www.crisprscan.org/?page=sequence) using the wildtype DNA sequence as input **(Figure 2a**). The user can also specify a preselected sgRNA spacer sequence from any other designing methods if desired. After validating the inputs, pegFinder first identifies the differences between the wildtype and edited DNA sequences by performing a Needleman-Wunsch alignment with affine gap penalties ^9^. Using the alignment, pegFinder chooses a single sgRNA spacer from the eligible candidate sgRNAs (**Figure 2b-c**). If a preselected sgRNA was specified, pegFinder will validate whether the chosen sgRNA is correctly positioned to produce the desired edits. pegFinder then identifies an appropriate RT template and PBS sequence to generate the desired edit by factoring in the positioning of the edited bases and the GC content of the sgRNA, respectively (**Figure 2d-e**). Additionally, pegFinder identifies secondary sgRNAs that nick 20-100nt away on the opposite strand from the primary sgRNA, which can potentially increase the efficiency of prime editing by favoring the edited strand during heteroduplex resolution (i.e. the PE3 method) ^2^ (**Figure 2f**). To facilitate rapid experimental implementation, pegFinder further generates oligonucleotide sequences that can be directly used to clone the pegRNAs into standard plasmid vectors (**Figure 2g, Methods**).

**Figure 1:**
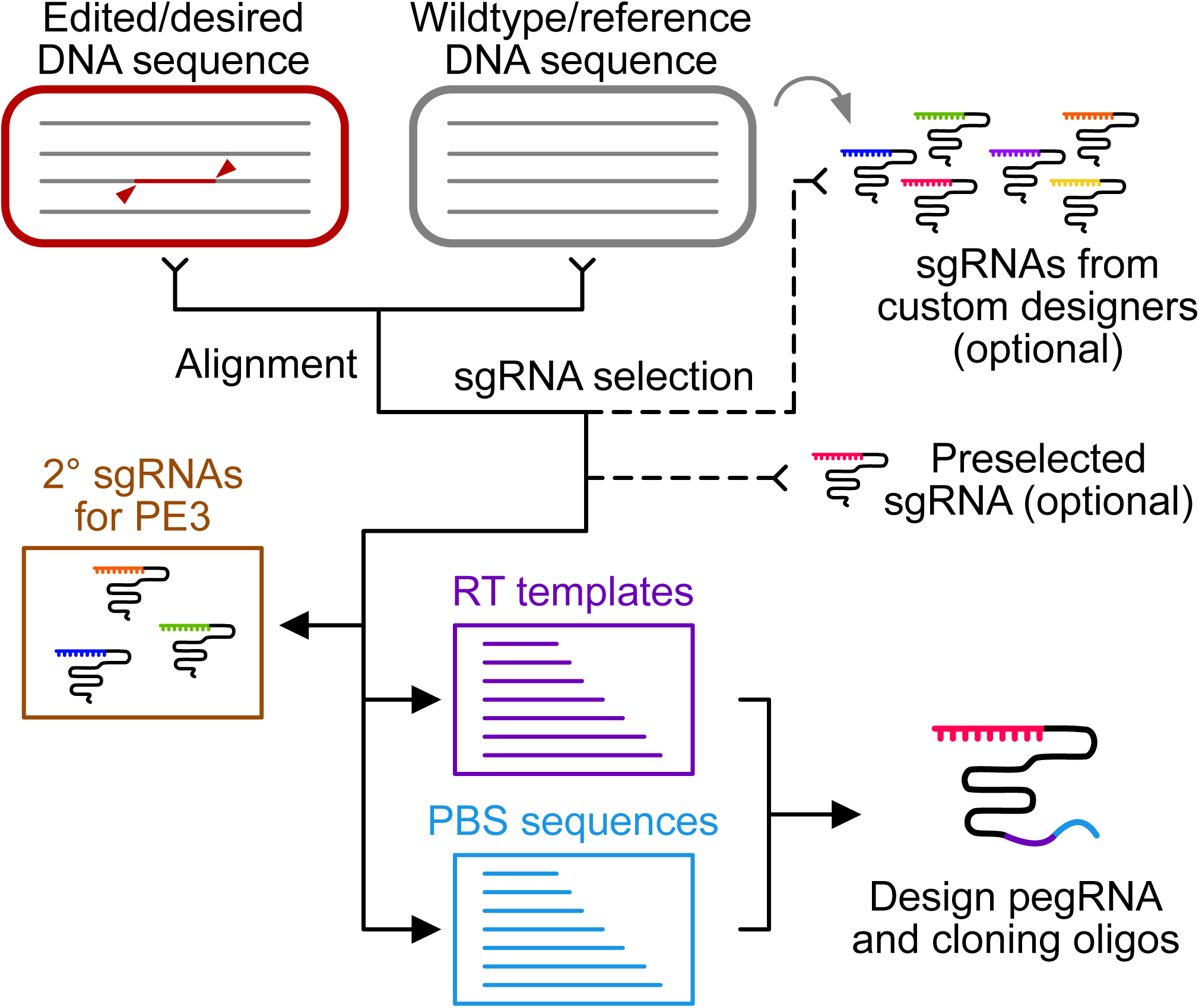
pegFinder: A pegRNA designer for CRISPR prime editing. Schematic of the pegFinder workflow for designing CRISPR prime editing pegRNAs. The user provides the wildtype DNA sequence, and the edited DNA sequence. Optionally, the user can include the results from sgRNA designer tools to pegFinder, or specify a preselected sgRNA spacer to be used for pegRNA design. pegFinder then generates a pegRNA that can be used to engineer the desired alterations. pegFinder also reports alternative RT templates and PBS sequences of varying lengths that can be swapped into the pegRNA for downstream experimental optimization. pegFinder further reports secondary nicking sgRNAs that can increase prime editing efficiency (PE3 method).

**Figure 2:**
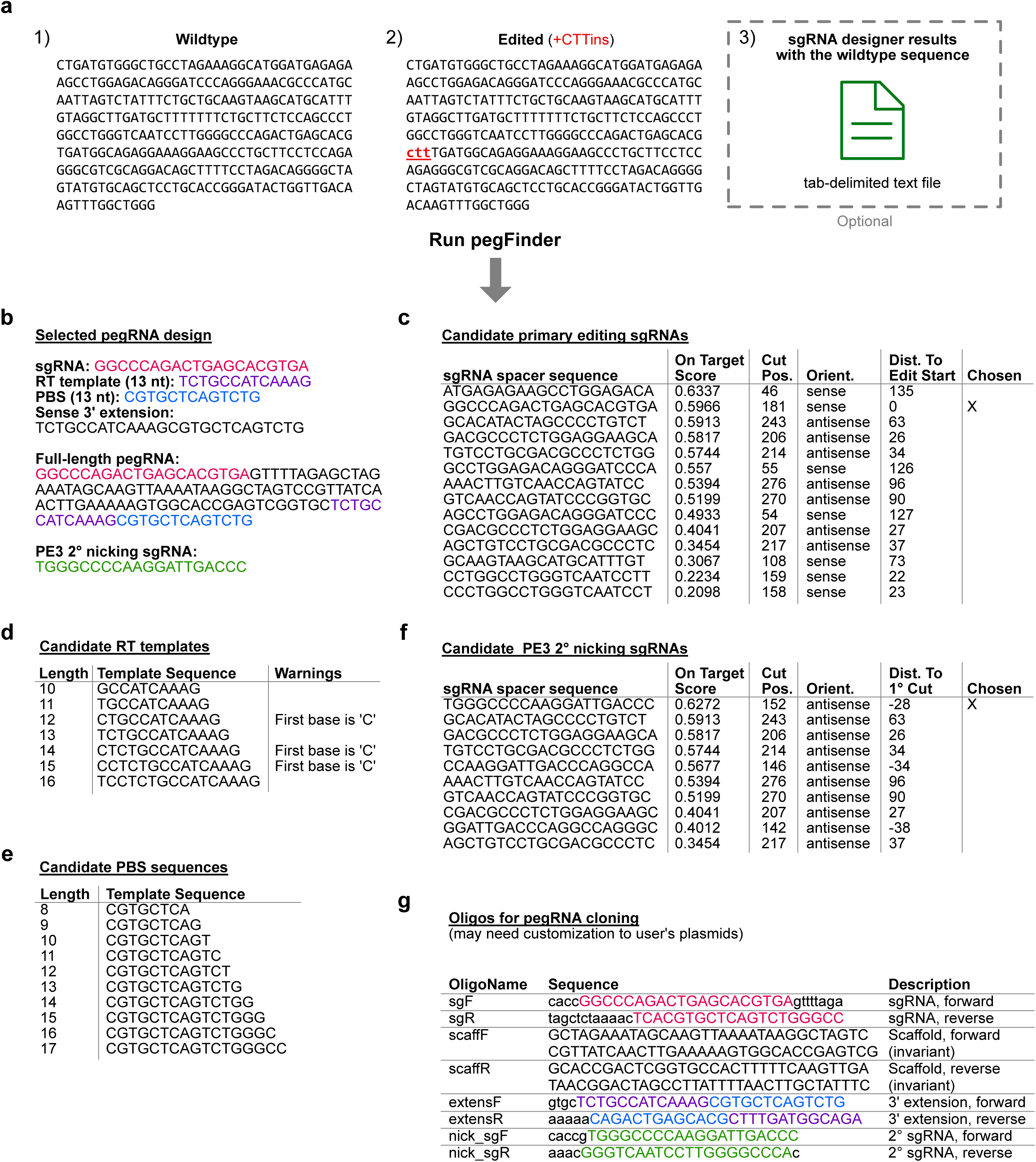
pegFinder workflow and example results. **a)** Example of pegFinder inputs: 1) wildtype DNA sequence, 2) edited DNA sequence showing a +CTT insertion, and 3) sgRNA designer results using the wildtype sequence as input (optional). **b)** pegFinder returns a pegRNA suitable for the desired edit, detailing the chosen sgRNA spacer, RT template, and PBS sequence, as well as the concatenated 3’ extension and full-length pegRNA sequence. pegFinder also reports a compatible secondary nicking sgRNA that can potentially increase prime editing efficiency. **c)** pegFinder identifies all candidate sgRNAs that could potentially mediate the desired edits. pegFinder chooses a single sgRNA by balancing on-target efficacy (if applicable) and distance to the edit start. **d)** pegFinder returns candidate RT templates of varying lengths that can be used for pegRNA optimization. **e)** pegFinder provides a list of candidate PBS sequences of varying lengths that can be used for pegRNA optimization. **f)** pegFinder reports all candidate PE3 secondary nicking sgRNAs that cut the opposite strand 20-100nt away from the primary sgRNA. From the candidates, pegFinder chooses the secondary nicking sgRNA with the highest on-target score (if applicable). **g)** pegFinder generates oligonucleotide sequences suitable for ligation cloning of the pegRNA and the secondary nicking sgRNA into standard expression vectors.

Given that prime editing has only recently been developed, the rules governing pegRNA design are incompletely understood. Thus, experimental optimization of the pegRNA for each given experimental application is likely necessary. Since the efficiency of prime editing is known to vary depending on the length and/or base composition of the RT template and PBS, pegFinder reports RT templates and PBS sequences of varying lengths (**Figure 2d-e**). pegFinder also returns all candidate primary and secondary sgRNA spacers to facilitate further downstream optimizations (**Figure 2c,f)**.

To experimentally validate the design algorithm, we used pegFinder to design two different pegRNAs targeting the human *HEK3* locus (also known as *LINC01509*). The first pegRNA was designed to insert “CTT” into the same genomic position as one of the pegRNAs described in the original prime editing study ^2^, and thus served as a positive control benchmark. With only minimal user inputs (**Figure 2a**), pegFinder designed a candidate CTTins pegRNA that was largely identical in sequence to the CTTins pegRNA described previously (Addgene #132778) ^2^, demonstrating the accuracy of the algorithm (**Figure 2b-e**). The oligonucleotide sequences generated by pegFinder (**Figure 2g**) were then directly used for ligation cloning (**Methods**). In comparison to control cells co-transfected with PE2 and empty vector (**Figure 3a**), cells co-transfected with PE2, the pegFinder-designed CTTins pegRNA showed evidence of prime editing, as determined by analysis of minor peaks in the sequencing chromatograms (**Figure 3b**). We similarly used pegFinder to design a pegRNA for inserting “CT” into the *HEK3* locus. Of note, this particular edit (CTins) was not performed in the original study ^2^. Using the constructs produced by pegFinder, we observed evidence of prime editing in cells transfected with the CTins pegRNA (**Figure 3c**), experimentally demonstrating the functionality of pegFinder-designed pegRNAs.

**Figure 3:**
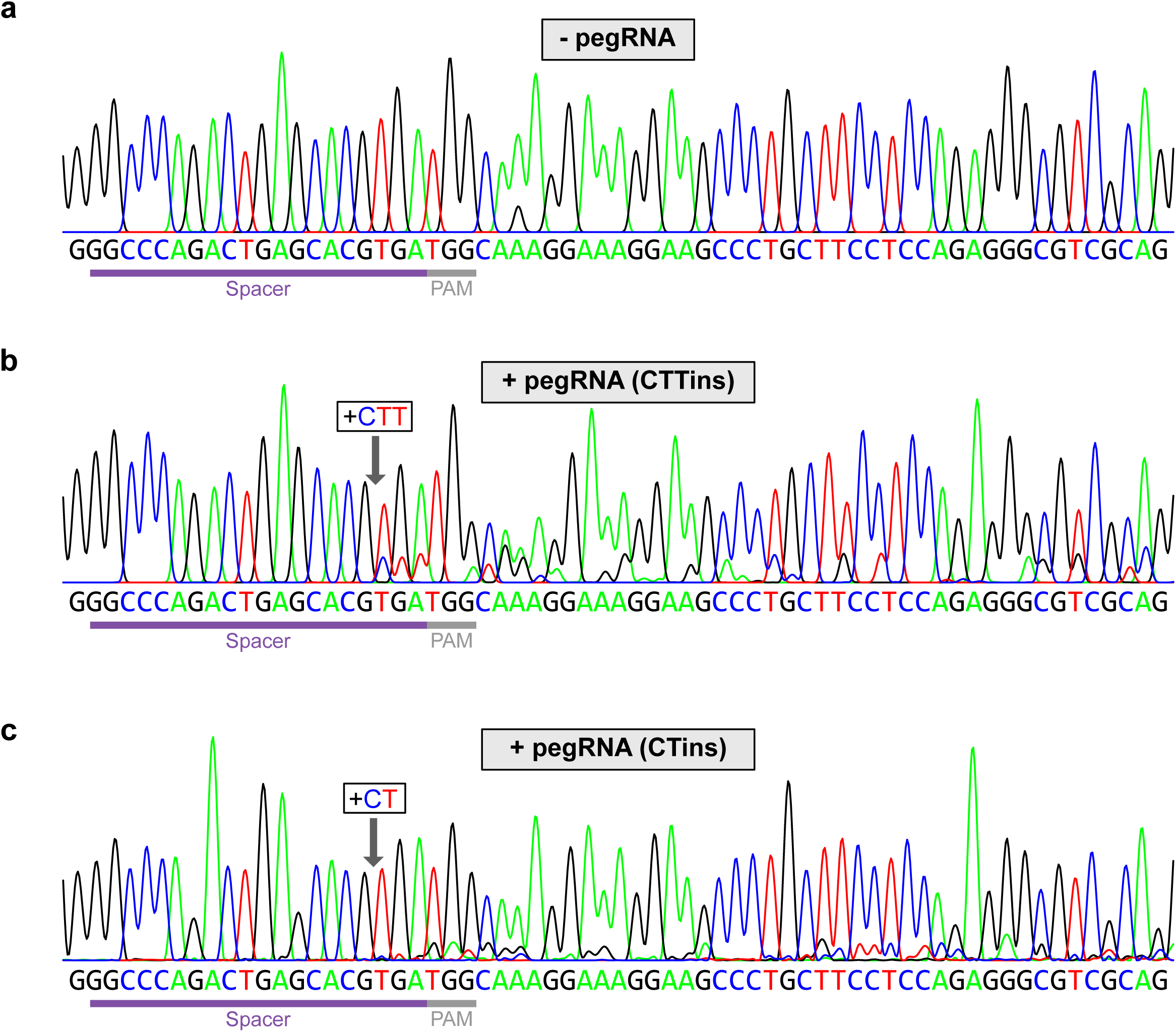
Experimental validation of pegRNAs designed by pegFinder. **a)** Sequencing chromatogram of the *HEK3* target site in cells transfected with vector control. **b)** Sequencing chromatogram of the *HEK3* target site in cells transfected with a pegRNA designed to insert “CTT” into the +1 position, along with a secondary nicking sgRNA. **c)** Sequencing chromatogram of the *HEK3* target site in cells transfected with a pegRNA designed to insert “CT” into the +1 position, along with a secondary nicking sgRNA.

Together, these data showcase the simplicity and utility of pegFinder for pegRNA design. pegFinder provides a convenient platform for researchers in diverse fields to rapidly harness the versatility of prime editing.

## Acknowledgments

We thank S. Eisenbarth for support. RDC is supported by the Yale MSTP training grant from NIH (T32GM007205) and an NIH NRSA fellowship from NCI (F30CA250249). JSC is supported by Yale MSTP training grant from NIH (T32GM007205). SC is supported by Yale SBI/Genetics Startup Fund, NIH/NCI/NIDA (DP2CA238295, 1R01CA231112, U54CA209992-8697, R33CA225498, RF1DA048811), AACR (499395, 17-20-01-CHEN), Ludwig Family Foundation, Sontag Foundation, Blavatnik Family Foundation and Chenevert Family Foundation.

## Author contributions

R.D.C. conceived the pegRNA design tool (pegFinder). R.D.C. developed the pegFinder algorithm. J.S.C. developed the web interface. R.D.C. and J.S. performed experiments. R.D.C. and J.S.C. wrote the manuscript. S.C. provided conceptual advice and supervised the work.

## Figure Legends

**Supplementary Figure 1:**
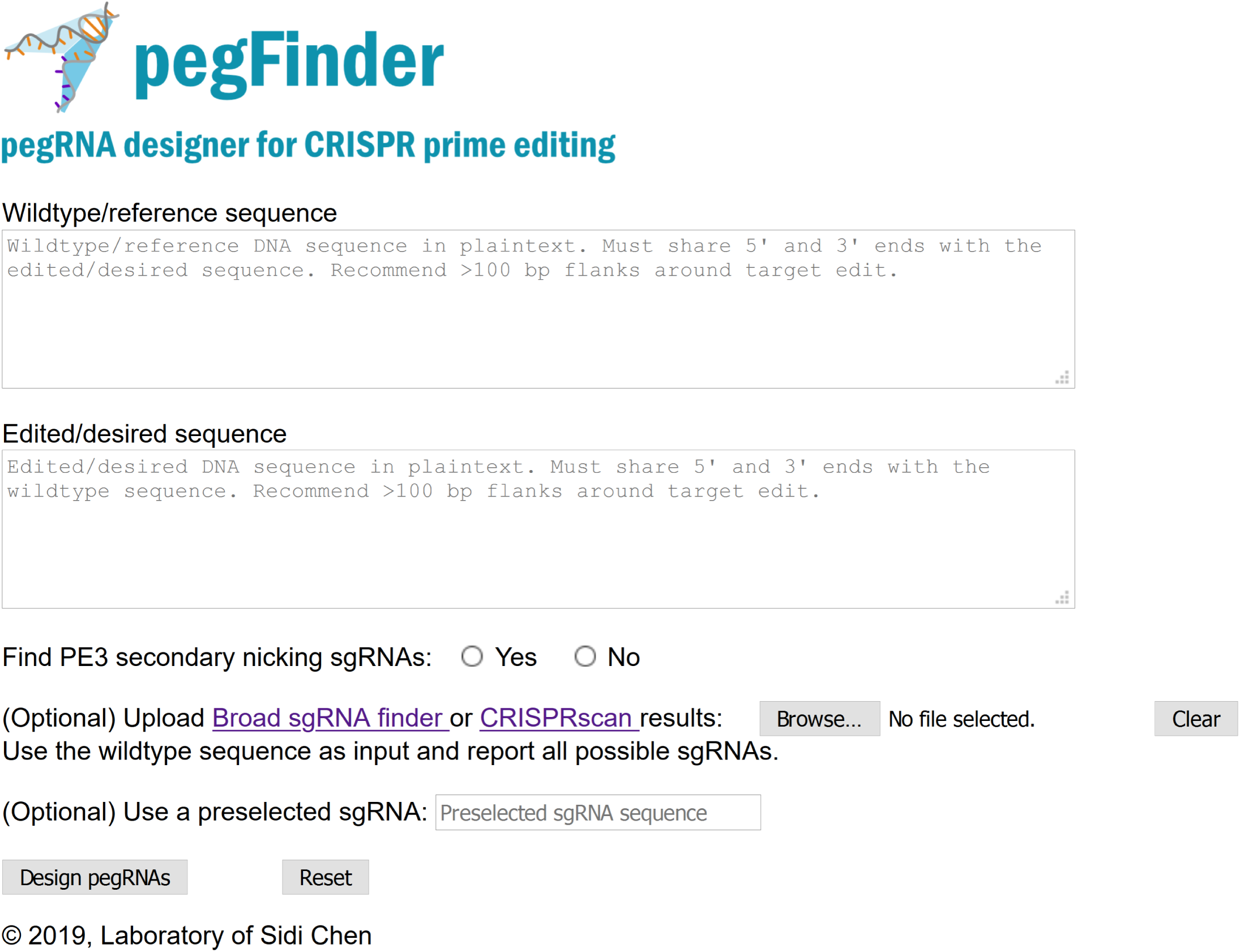
pegFinder web interface. Screenshot of the pegFinder web interface, available at http://pegfinder.sidichenlab.org.

## Supplementary Tables

**Table S1:** Input DNA sequences used in this study for experimental validation of pegFinder-designed pegRNAs.

**Table S2:** Example of Broad sgRNA designer results, using the wildtype *HEK3* target sequence in Table S1 as input.

**Table S3:** Example of CRISPRscan sgRNA designer results, using the wildtype *HEK3* target sequence in Table S1 as input.

## Methods

### Development of pegFinder algorithm and web server

The pegFinder core algorithm was developed in Perl. The web portal was implemented in the Mojolicious - Perl real-time web framework.

### Inputs for pegFinder

pegFinder minimally requires two inputs. First, the wildtype DNA sequence of the region of interest is needed. Since candidate sgRNAs must be found within this wildtype sequence, we recommend >100nt flanks around the desired edit site. These sequences can be readily retrieved through genome browsers such as UCSC, IGV, or Ensembl BioMart. Second, the edited DNA sequence should be obtained by modifying the wildtype sequence to incorporate the desired alterations. Note that pegFinder expects the wildtype and edited sequences to share identical 5’ and 3’ ends, and will notify the user when this is not the case. As an example, consider a 200nt wildtype DNA sequence in which the user wishes to insert a 10nt sequence after the 100^th^ nucleotide. The edited sequence should then be 100nt flank – 10nt insertion – 100nt flank, where the 5’ and 3’ 100nt flanks around the insert correspond exactly to the 200nt wildtype sequence. Thus, we recommend generating the edited DNA sequence by directly modifying the wildtype DNA input, as doing so ensures that the 5’ and 3’ flanks will remain identical.

Optionally, pegFinder can incorporate predicted on-target/off-target scores from sgRNA designer tools, with the caveat that these scoring algorithms were trained on gene knockout data, and thus may not be relevant for prime editing experiments. pegFinder can use the results from the Broad sgRNA designer tool^5^ (https://portals.broadinstitute.org/gpp/public/analysis-tools/sgrna-design) or CRISPRscan^4^ (https://www.crisprscan.org/?page=sequence), with the wildtype DNA sequence described above as the input query. For the Broad designer, the CRISPR enzyme should be selected as SpyoCas9 (NGG), and the appropriate target genome should be selected. Note also that all unpicked sequences should be reported. The tab-delimited results file that is produced (“sgRNA Picking Results”) can be saved and uploaded to pegFinder. For CRISPRscan, the correct target species should be chosen, the enzyme should be set as Cas9-NGG, and the option to find sgRNAs from both T7 and Sp6 promoters should be selected. The tab-delimited file that is produced can be saved and uploaded to pegFinder. As another alternative, if the user wishes to specify a preselected sgRNA, the 20nt spacer sequence can be entered directly into pegFinder.

### Alignment of wildtype and edited DNA sequences

The first step of pegFinder is to align the wildtype and edited DNA sequences using the Needleman-Wunsch algorithm, with an affine gap penalty function ^9^. The positions of gaps and mismatched bases are noted, which are then used to select candidate sgRNAs.

### Selection of primary nicking sgRNA spacers for pegRNAs

If no preselected sgRNA spacer was specified, pegFinder can either identify candidate sgRNA spacers *de novo* or utilize the outputs from sgRNA designer tools. After determining the position range of the desired alterations, pegFinder searches for spacers that could potentially mediate the desired prime editing outcome. pegFinder designates sgRNA spacers on the “sense” strand as potential candidates if these conditions are met: 1) the sgRNA cut position is upstream of the 5’-most edited base, 2) the 3’-most edited base is within 150nt of the cut position, and if applicable, 3) the sgRNA spacer has a total “off-target” Tier I Bin I + Bin II + Bin III score ≤ 1 (as determined by the Broad sgRNA designer), or the number of “seed off-targets” is 0 (as determined by CRISPRscan). While a Tier I score of 1 on the Broad designer would normally indicate a perfect-match off-target site, in this context it would simply correspond to the genomic locus of the input sequence, and thus can be ignored. Similarly, for sgRNA spacers on the “antisense” strand, pegFinder designates a spacer as a candidate if: 1) the sgRNA cut position is downstream of the 3’-most edited base (in the sense orientation), 2) the 5’-most edited base (sense orientation) is within 150nt of the cut position, and if applicable, 3) the sgRNA has a total “off-target” Tier I Bin I + Bin II + Bin III score ≤ 1 (Broad sgRNA designer) or a “seed off-targets” count of 0 (CRISPRscan).

In standalone mode, pegFinder then selects the candidate sgRNA spacer that is closest to the first edited base. When incorporating on-target prediction scores from sgRNA designer tools, pegFinder selects a single sgRNA spacer by balancing predicted on-target efficacy with distance to the edit start. If there are candidate spacers with on-target scores ≥ 0.5 (Broad designer) or ≥ 25 (CRISPRscan), pegFinder will choose the sgRNA with the shortest distance to the closest edit position. In the event of a tie for shortest distance, pegFinder chooses the sgRNA with the higher on-target score. If there no candidate sgRNAs with high on-target scores, pegFinder will then choose the sgRNA with the highest on-target score regardless of its distance to the edit start (up to 150nt). While pegFinder automatically chooses a single sgRNA spacer for further pegRNA design, all candidate sgRNA spacers are reported in the pegFinder outputs. If, for instance, subsequent experiments demonstrate that the sgRNA spacer originally chosen by pegFinder is inefficient at inducing prime editing, the user can easily rerun pegFinder by prespecifying a different sgRNA spacer from the list of potential candidates.

### Selection of secondary nicking sgRNA spacers for PE3

Nicking the opposite strand can increase the efficiency of prime editing (an approach termed PE3) ^2^. To design “secondary” nicking sgRNAs to be used for PE3, pegFinder can either identify sgRNA spacers *de novo* or utilize the results from sgRNA designer tools (Broad sgRNA designer or CRISPRscan) with the wildtype DNA sequence as input. pegFinder considers an sgRNA to be a candidate for secondary nicking if: 1) the sgRNA targets the strand opposite of the primary nicking sgRNA, 2) the secondary nick occurs 20-100nt away from the primary nick, and if applicable, 3) the sgRNA has a sum “off-target” Tier I Bin I + Bin II + Bin III score ≤ 1 (Broad designer) or the number of “seed off-targets” is 0 (CRISPRscan). In standalone mode, pegFinder identifies the sgRNA spacer that nicks closest to ± 50nt from the primary nicking sgRNA. When incorporating on-target efficacy predictions, pegFinder chooses the sgRNA spacer with the highest on-target score among the candidate secondary nicking sgRNAs. All candidate secondary nicking sgRNA spacers are returned by pegFinder for ease of experimental optimization.

### Selection of RT templates and PBS sequences for comprising pegRNAs

After selecting a primary sgRNA (or directly using the user-specified preselected sgRNA), pegFinder then extracts candidate RT templates and PBS sequences that can be incorporated into the 3’ extension of pegRNAs. For designing RT templates, pegFinder uses the edited/desired sequence and extracts the DNA between the primary nick site and the farthest edited base (the 3’-most base if using a sense strand sgRNA, or the 5’-most base if antisense), plus an additional 1nt. If the resultant sequence is < 10nt, pegFinder will report candidate RT templates ranging from 10-16nt in length. If the distance between the primary nick site and the farthest edited base is ≥ 10nt, pegFinder will report RT templates ranging from +1 to +7nt of the nick to edit distance. In all cases, pegFinder will flag RT templates that have a “C” as their 5’ most base (corresponding to a “G” as the final templated base), since it was previously demonstrated that such RT templates exhibit lower efficiency for prime editing, potentially due to basepairing interactions with the sgRNA scaffold ^2^. By default, pegFinder will then select a single RT template by choosing the template representing the median length among the candidates that do not begin with “C”, choosing the shorter template if there are an even number of candidates. If no RT templates exist that do not begin with “C”, pegFinder will select the template of median length among all candidates. Since the length of the RT template may require further optimization, all candidate RT templates are also reported by pegFinder.

To design PBS sequences, pegFinder extracts sequences 8-17nt in length from the reverse complement of the primary sgRNA sequence, moving backwards from the −1 position (the position before the cut site). If the GC% of the primary sgRNA is between 30-70%, pegFinder will choose the PBS sequence of 13nt by default. If GC% < 30%, pegFinder chooses the PBS with length 14nt, and if GC% > 70%, pegFinder chooses the PBS of length 12nt. All PBS sequences 8-17nt are outputted by pegFinder to facilitate experimental optimization.

### Design of oligonucleotide sequences for cloning pegRNAs and sgRNAs

After choosing a primary sgRNA spacer, secondary sgRNA spacer, RT template, and PBS sequence, pegFinder additionally outputs oligonucleotide sequences that can be directly utilized for ligation cloning of the designed pegRNAs and/or sgRNAs. The oligos designed by pegFinder are intended for ligation cloning through an adaption of the lentiGuide-Puro protocol ^10^ (see below). Of note, pegRNA/sgRNA sequences that do not begin with a “G” on the 5’ end will automatically have a “G” appended in the cloning oligonucleotide sequences to facilitate transcription from the standard U6 promoter.

### pegRNA/sgRNA cloning protocol using oligos designed by pegFinder

Using the oligonucleotide sequences produced by pegFinder, each forward/reverse oligo pair was annealed in T4 Ligation Buffer (NEB), with T4 PNK to phosphorylate the oligos. The recipient pegRNA expression vector (pU6-pegRNA-GG-acceptor vector; Addgene #132777) was digested by BsaI (NEB). After diluting the oligo duplexes 1:100, the primary nicking sgRNA, the invariant scaffold, and the 3’ extension were ligated together into the digested vector using Quick Ligase or T4 Ligase (NEB) to generate the complete pegRNA plasmid. Similarly, the secondary nicking sgRNA diluted duplex was ligated into a standard U6 sgRNA vector (such as the lentiGuide-Puro vector; Addgene #52963). Note that when using alternative plasmids for performing pegRNA/sgRNA cloning, the overhangs produced by pegFinder may need to be customized to match the sticky ends following plasmid digestion.

### Experimental validation of pegRNAs

For the +1 CTT insertion at *HEK3*, pegFinder designed the following pegRNA: 5’ GGCCCAGACTGAGCACGTGAGTTTTAGAGCTAGAAATAGCAAGTTAAAATAAGGCTAGTCCGTTATCAACTTGAAAAAGTGGCACCGAGTCGGTGCTCTGCCATCAAAGCGTGCTCAGTCTG 3’.

For the +1 CT insertion at *HEK3*, pegFinder designed the following pegRNA: 5’ GGCCCAGACTGAGCACGTGAGTTTTAGAGCTAGAAATAGCAAGTTAAAATAAGGCTAGTCCGTTATCAACTTGAAAAAGTGGCACCGAGTCGGTGCTCTGCCATCAAGCGTGCTCAGTCTG 3’.

The oligo sequences provided by pegFinder were then cloned into the appropriate expression vectors, as described above. Experimental validation of the pegRNAs was performed as described previously, with minor modifications ^2^. HEK293T cells (ATCC) were seeded on 24-well plates and transfected 16 hours later at 60-80% confluency with 2 ul Lipofectamine 2000 (ThermoFisher), 1.5 ug pCMV-PE2 plasmid (Addgene #132775), 500 ng pegRNA plasmid (cloned into Addgene #132777), and 200 ng secondary nicking sgRNA plasmid. 50 ng sfGFP-N1 (Addgene #54737) was also included to assess transfection efficiency. Cells were harvested 2 days post-transfection and genomic DNA (gDNA) was purified with the QIAamp DNA Blood Mini Kit (Qiagen). The genomic region surrounding the pegRNA target site was then amplified by PCR with the following primers, using 200 ng input gDNA: Forward, AGGGAAACGCCCATGCAATTAGTCT Reverse, CTAGCCCCTGTCTAGGAAAAGCTGTC

PCR was performed using Phusion Flash High-Fidelity polymerase (ThermoFisher) with the following settings: 98°C for 2 min, then 35 cycles of [98°C for 1 s, 60°C for 5 s, and 72°C for 5 s], followed by 72°C extension for 2 min. The resulting PCR amplicons were gel-purified (Qiagen) and processed for Sanger sequencing (Applied Biosystems 3730xL DNA Analyzer).

## Data availability statement

For the pegRNAs that were experimentally tested in this study, all relevant information is provided in the Supplementary Tables. This information can be used to recreate the pegRNA designs described here, through the pegFinder web portal available at http://pegfinder.sidichenlab.org.

## Code availability statement

The code that supports the findings of this study is available at GitHub (https://github.com/rdchow/pegfinder). The web portal server is accessible at http://pegfinder.sidichenlab.org for non-profit use.

## References

1. Pickar-Oliver, A. & Gersbach, C. A. The next generation of CRISPR–Cas technologies and applications. Nat Rev Mol Cell Biol 20, 490–507 (2019).

2. Anzalone, A. V. et al. Search-and-replace genome editing without double-strand breaks or donor DNA. Nature 576, 149–157 (2019).

3. Meier, J. A., Zhang, F. & Sanjana, N. E. GUIDES: sgRNA design for loss-of-function screens. Nature Methods 14, 831–832 (2017).

4. Moreno-Mateos, M. A. et al. CRISPRscan: designing highly efficient sgRNAs for CRISPR-Cas9 targeting in vivo. Nat. Methods 12, 982–988 (2015).

5. Doench, J. G. et al. Optimized sgRNA design to maximize activity and minimize off-target effects of CRISPR-Cas9. Nat. Biotechnol. 34, 184–191 (2016).

6. Haeussler, M. et al. Evaluation of off-target and on-target scoring algorithms and integration into the guide RNA selection tool CRISPOR. Genome Biology 17, 148 (2016).

7. Labun, K. et al. CHOPCHOP v3: expanding the CRISPR web toolbox beyond genome editing. Nucleic Acids Res 47, W171–W174 (2019).

8. Park, J., Bae, S. & Kim, J.-S. Cas-Designer: a web-based tool for choice of CRISPR-Cas9 target sites. Bioinformatics 31, 4014–4016 (2015).

9. Needleman, S. B. & Wunsch, C. D. A general method applicable to the search for similarities in the amino acid sequence of two proteins. Journal of Molecular Biology 48, 443–453 (1970).

10. Sanjana, N. E., Shalem, O. & Zhang, F. Improved vectors and genome-wide libraries for CRISPR screening. Nat. Methods 11, 783–784 (2014).

